# Convergence of virulence and multidrug resistance in a single plasmid vector in multidrug-resistant *Klebsiella pneumoniae* ST15

**DOI:** 10.1101/463406

**Authors:** Margaret M. C. Lam, Kelly L. Wyres, Ryan R. Wick, Louise M. Judd, Aasmund Fostervold, Kathryn E. Holt, Iren Høyland Löhr

## Abstract

**Background:** Multidrug resistance (MDR) and hypervirulence (hv) are typically observed in separate *Klebsiella pneumoniae* populations. However, convergent strains with both properties have been documented and potentially pose a high risk to public health in the form of invasive infections with limited treatment options.

**Objectives:** To characterize the genetic determinants of virulence and antimicrobial resistance (AMR) in two ESBL-producing *K. pneumoniae* isolates belonging to the international MDR clone ST15.

**Methods:** The complete genome sequences of both isolates, including their plasmids, were resolved using Illumina and Oxford Nanopore sequencing.

**Results:** Both isolates carried large mosaic plasmids in which AMR and virulence loci have converged within the same vector. These closely related mosaic hv-MDR plasmids include sequences typical of the *K. pneumoniae* virulence plasmid 1 (KpVP-1; including aerobactin synthesis locus *iuc*) fused with sequences typical of IncFII_K_ conjugative AMR plasmids. One hv-MDR plasmid carried three MDR elements encoding the ESBL gene *bla*_CTX-M-15_ and eight other AMR genes (*bla*_TEM_, *aac3’-IIa*, *aph3’-Ia, dfrA1*, *satA2*, *bla*_SHV_, *sul1*, *aadA1*). The other carried remnants of these elements encoding *bla*_TEM_ and *aac3’-IIa*, and *bla*_CTX-M-15_ was located in a second plasmid in this isolate. The two isolates originated from patients hospitalized in Norway but have epidemiological and genomic links to Romania.

**Conclusions:** The presence of both virulence and AMR determinants on a single vector enables simultaneous transfer in a single event and potentially rapid emergence of hv-MDR *K. pneumoniae* clones. This highlights the importance of monitoring for such convergence events with stringent genomic surveillance.

## INTRODUCTION

The majority of infections caused by *Klebsiella pneumoniae* (*Kp*) are typically associated with one of two distinct clinical phenomena caused by non-overlapping *Kp* populations: healthcare-associated infections caused by MDR *Kp* strains that also often cause nosocomial outbreaks, and community-acquired, invasive infections caused by hypervirulent (hv) strains ^1,2^. However, convergent strains carrying both MDR and hypervirulent genes have been reported ^3–7^. Recently, a high-mortality outbreak of ventilator-associated pneumonia caused by a strain of hv carbapenemase-producing sequence type (ST) 11 *Kp* was reported in China, demonstrating that the combination of enhanced virulence potential and difficulties in treatment posed by MDR can be fatal. The Chinese report was particularly notable as ST11 is typically associated with MDR, and appears to be the most common cause of carbapenemase-producing *Kp* infections reported in China. However, the outbreak strains had additionally acquired a virulence plasmid harbouring *iuc* (aerobactin siderophore) and *rmpA2* (hypermucoidy) loci, which are usually only observed in hypervirulent clones, such as ST23 ^2,8,9^.

Given that antimicrobial resistance (AMR) and virulence determinants are commonly mobilized on plasmids, their occasional convergence within individual strains is not unexpected. The highly mosaic nature of *Kp* plasmids creates the risk of AMR and virulence determinants converging within a single plasmid. This hv-AMR vector could spread amongst *Kp* and confer widespread ability to cause serious infections with very limited treatment options. To our knowledge only two such plasmids have been reported; pKpvST147L, harbouring *iuc, rmpA, rmpA2* and several AMR determinants (*sul2, armA, sul1* and *mphA*) in a ST147 carriage isolate also carrying a *bla*_NDM-1_ carbapenemase and isolated in London ^4^; and pKP70-2, harbouring the typical KpVP1 virulence plasmid of ST23 (encoding *iuc, iro*, *rmpA* and *rmpA2*) with an additional insertion of a MDR transposon including *bla*_KPC-2_ carbapenemase in a K1 ST23 sputum isolate isolated in China ^10^.

Here we report the complete genome sequences of two *Kp* ST15 carrying both MDR and virulence determinants, identified during a study of ESBL-producing *Kp* isolates from Norwegian hospitals.

## MATERIALS AND METHODS

### Ethics statement

The isolates presented here were collected and sequenced as part of a larger national study of *Kp* in Norwegian hospitals between 2001 and 2015 called NOR-KLEB. Ethical approval for NOR-KLEB, including the collection and sequencing of *Kp* isolates and collection of patient data, was provided by the Regional ethics committee: REC west, application ID:2017/1185.

### Bacterial isolates

Isolate KP_NORM_BLD_2014_104014 (KP_104014) was cultured from an 86-year-old Romanian male admitted to an Oslo hospital in 2014 with cholangiocarcinoma before developing bacteremia. Isolate KP_NORM_BLD_2015_112126 (KP_112126) was cultured from a 76-year-old female admitted to a Western Norway hospital in 2015 to treat a glioblastoma, and developed neutropenic fever with pneumonia and bacteremia. She had been hospitalized in Romania prior to admission in Norway. Antimicrobial susceptibility was determined by disk diffusion and broth micro dilution, and hypermucoidy was assessed via the string test.

### Whole genome sequencing and analysis

250 and 150 bp paired end reads were generated for n=12 ST15 Norwegian *K. pneumoniae* isolates on the Illumina MiSeq and HiSeq platforms, respectively, and assembled with Unicycler v0.4.4-beta. In order to resolve the complete plasmid sequences for strains KP_104014 and KP_112126, additional long read sequencing on a MinION R9.4 flow cell (Oxford Nanopore Technologies) was performed, and combined with the Illumina short reads to generate hybrid assemblies, using Unicycler as previously described ^11,12^, which were annotated using Prokka v1.11 ^13^. Genotyping information including multi-locus sequence type (MLST), capsule type, AMR and virulence gene detection was extracted using Kleborate (https://github.com/katholt/Kleborate) and used to curate the annotation of relevant loci in the plasmids.

To place the hv-MDR strains in context, we performed comparative genomic analyses (described below) with an additional n=10 ST15 strains isolated between 2003 and 2015 from seven hospitals across Norway as part of the NOR-KLEB study (full results to be reported elsewhere), together with publicly available Illumina data identified from papers reporting *Kp* ST15 genome sequences (genomes and references listed in **Supplementary data 1**). Illumina read data for *Kp* genomes collected by the EuSCAPE European survey of carbapenemase-producing Enterobacteriaceae ^20^ were downloaded and assembled using Unicycler and genotyped using Kleborate to identify ST15 isolates, and the ST15 read sets were included in the comparative analysis.

All read sets were mapped to the genome of KP_104014 using the RedDog v1b 10.2 pipeline (https://github.com/katholt/RedDog). An alignment of chromosomal single nucleotide variants was extracted, recombinant regions were identified and filtered from the alignment using Gubbins v2.0.0 ^14^, and the final alignment was passed to RAxML v8.1.23 ^15^ to infer a core genome maximum-likelihood phylogeny. From the mapping data we also extracted the coverage of pKp104014_1 sequence, the coverage of the hv-MDR plasmid of KP_104014, and the presence of genes annotated in pKp104014_1 (presence defined as ≥95% of the length of the gene covered by ≥five reads).

### Nucleotide Data Accessions

Complete, annotated sequences for the two novel genomes have been deposited in FigShare (doi:10.6084/m9.figshare.7222889) and GenBank (BioSamples SAMEA5063299 and SAMEA5063300). The accessions for the mosaic plasmids described here are TBC (pKp104014_1) and TBC (pKp112126_1).

## RESULTS AND DISCUSSION

Isolate KP_104014 displayed resistance to cefotaxime, ceftazidime, ciprofloxacin, gentamicin, piperacillin-tazobactam and co-trimoxazole, and susceptibility to meropenem, colistin and tigecycline. The complete genome sequence resolved seven plasmids (**Supplementary data 2**), including a novel 346 kbp mosaic hv-MDR plasmid pKp104014_1, which harboured *bla*_CTX-M-15_ and seven additional AMR genes (**Figure 1**). Plasmid pKp104014_1 shares regions of homology with typical KpVP-1- type *Kp* IncFIB_K_ virulence plasmids ^9^, such as pK2044 (40% coverage, including *iuc* and *rmpA2*), in addition to regions of homology to IncFII_K_ conjugative AMR plasmids (closest match: 246 kbp plasmid pKp_Goe_579-1, accession CP018313.1, from a ST147 *Kp* isolated in Germany, 59% coverage). The IncFII_K_ regions include genes for conjugative transfer, suggesting the plasmid may be self-transmissible. The plasmid harboured eight AMR genes mobilized via various elements including *bla*_CTX-M-15_ (mobilized by IS*EcP1*); *bla*_TEM-1_ and *aac3’-IIa* (Tn*3*); *dfrA1* and *sat2* (In*2*/Tn*7*); *bla*_SHV-5_ (IS*26*); and *sul1* and *aadA1* (In*1*/Tn*21*) (**Figure 1**). A second copy of *bla*_CTX-M-15_ was also inserted into the chromosomal gene *phoE* (via *ISEcp1*), and additional AMR genes (*aacA4, bla_OXA-1_*, *bla_TEM-1_,* and *cat*) were carried on a 76 kbp IncFII plasmid, pKp104014_3 (**Supplementary data 2**).

**Figure 1.**
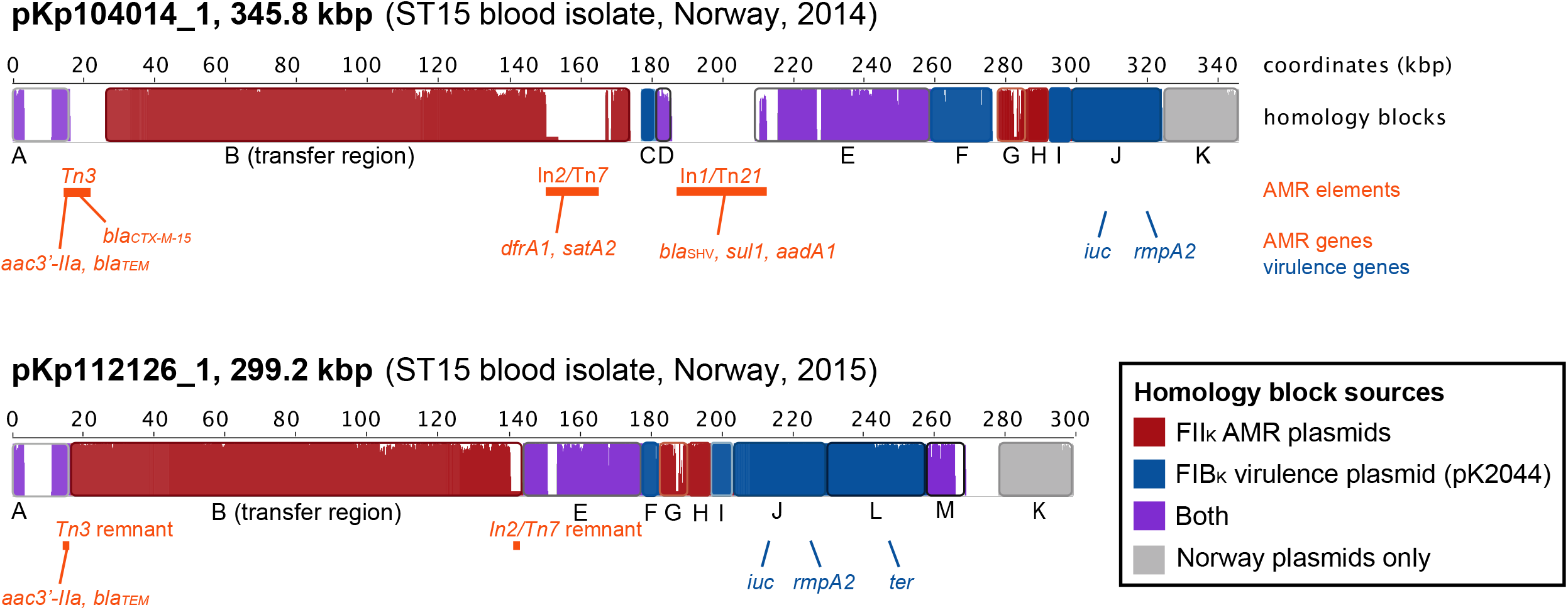
Map of novel mosaic hv-MDR plasmids, showing regions of homology with closely related AMR (pKp_Goe_579-1) and virulence (pK2044) plasmids, generated using Mauve. The location of known virulence genes (blue), as well as AMR genes and their associated mobile elements (red), are also indicated.

Isolate KP_112126 displayed resistance to cefotaxime, ceftazidime, ciprofloxacin and gentamicin, intermediate susceptibility piperacillin-tazobactam and tigecycline, and susceptibility to meropenem, colistin and co-trimoxazole. The complete genome sequence resolved four plasmids, including a mosaic 299 kbp hv-MDR plasmid pKp112126_1, with similarity to pKp104014_1 (99.99% nucleotide identity) including *iuc*, *rmpA2*, and the IncFII_K_ transfer region. This plasmid lacks most of the AMR genes, although remnants of two AMR regions of pKp104014_1, including one end of Tn*3* (encoding *bla*_TEM_, *aac3’-IIa*) and one end of In*2*/Tn*7* (integrase only), were present (**Figure 1**). Plasmid pKp112126_1 also carried an additional region with homology to KpVP1 virulence plasmids, including the *ter* locus encoding tellurite resistance (block L in **Figure 1**). *bla*_CTX-M-15_ was present in a distinct 90 kbp plasmid, pKp112126_3, which displayed homology with pKp104014_3 and *Shigella flexneri* plasmid R100 (accession AP000342.1).

Both of the Norwegian hv-MDR isolates belonged to ST15 and carried the siderophore yersiniabactin (in genomic island ICE*Kp2*) and the KL24 locus encoding capsular serotype K24. ST15 is a well-documented international ESBL-producing clone associated with nosocomial outbreaks worldwide, which frequently carries *bla*_CTX-M-15_-encoding IncFII plasmids ^16–19^. To explore the relatedness of the Norwegian isolates to one another and to the wider ST15 *Kp* population, we constructed a recombination-filtered, core-genome maximum likelihood phylogeny including KP_104014, KP_112126, ten additional ST15 isolates from Norway and 306 publicly available ST15 genomes from 29 other countries (**Figure 2, Supplementary data 1**). The tree showed that the two Norwegian hv-MDR isolates were closely related to one another (77 SNPs, 0.001% nucleotide divergence) and a urine isolate from Romania collected in 2013 (110 SNPs), but quite distant (>0.003% divergent) from the other Norwegian and global isolates.

**Figure 2.**
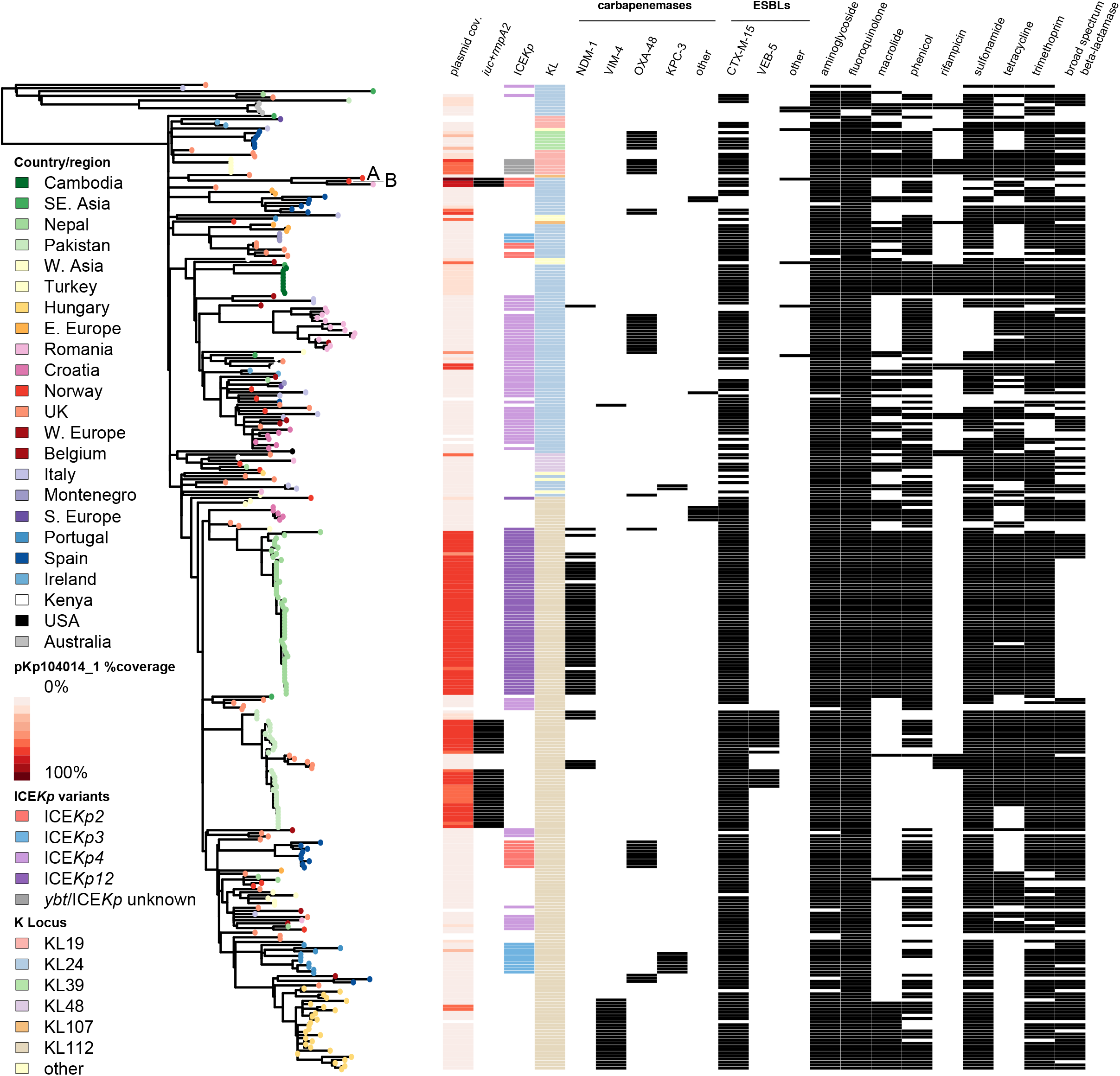
Recombination-free maximum likelihood phylogeny showing virulence and antimicrobial resistance properties of 152 ST15 isolates. Tips are coloured by the country of isolation (for n≥4 genomes) or geographical region as indicated, Norwegian convergent hv-MDR strains KP_NORM_BLD_2014_104014 and KP_NORM_BLD_2015_112126 labelled as A and B respectively. Columns are as follows: (1) % Coverage of the *bla*_CTX-M-15_/*iuc* plasmid pKp104014_1, determined by read mapping; (2) presence of the aerobactin synthesis locus *iuc* and the hypermucoidy *rmpA2* gene (coloured in black); (3) yersiniabactin ICE*Kp* variants detected; (4) capsule K-locus (KL); followed by antimicrobial resistance determinants as labelled (coloured in black for presence).

Interestingly, both the Norwegian hv-MDR plasmids were isolated from patients with epidemiological links to Romania (one of Romanian descent, one with recent travel history to Romania), suggesting the convergence of AMR and virulence plasmids may have occurred in that country rather than in Norway. The closely related Romanian isolate genome (ENA accession ERR1415588) carried *iuc* and *rmpA2* and its reads covered 98% of the pKp112126_1 sequence and only 54% of the typical virulence plasmid pK2044. This is consistent with the presence of a mosaic plasmid in this isolate, although the available Illumina reads were not sufficient to resolve the full sequence of the Romanian plasmid containing *iuc.*

*bla*_CTX-M-15_ was present in most (87%) of the ST15 genomes, along with other AMR genes (see **Figure 2, Supplementary data 1**). There were also multiple independent acquisitions of the ICE*Kp* genomic island encoding yersiniabactin, affecting 48% of all ST15 isolates including 50% of ESBL isolates (**Figure 2**). The only non-Norwegian ST15 isolates harbouring *iuc* were 30 isolates from Pakistan and the closely related Romanian isolate, all of which carried *iuc* and *rmpA2* loci in addition to *bla*_CTX-M-15_ and multiple other AMR genes. The convergence of AMR and virulence was noted in the original study reporting these genomes from Pakistan ^21^, however it is not possible to determine from the draft genomes whether *iuc* is co-localised on the same plasmid as AMR genes. Mapping of all ST15 read sets to pKp104014_1 showed that *iuc*+ isolates from Pakistan and *iuc*- isolates from Nepal (alongside a small number of *iuc-* isolates from other countries) share many genes with the mosaic plasmid pKp104014_1 (55.7-68.5% coverage for Pakistan isolates; 52.2-70.2% coverage for Nepal isolates) (**Figure 2** and **Supplementary figure**). This confirms that IncFII_K_ and IncFIB_K_ AMR and virulence plasmids circulate in South Asian *Kp* ST15 populations and could potentially fuse to form hybrid hv-MDR plasmids.

Concerningly, our findings reveal mosaic plasmids carrying both virulence determinants (*iuc, rmpA2*) and AMR determinants in ESBL-producing isolates of a well-established MDR *Kp* clone that has been associated with nosocomial infections and outbreaks worldwide. The co-presence of these loci in a single plasmid vector poses a substantial public health threat with the ability to simultaneously spread AMR and virulence, and it highlights the need for surveillance of virulence alongside AMR before such strains become widespread.

## Acknowledgments

We thank The Norwegian Surveillance System for Antimicrobial Drug Resistance (NORM) for data sharing and The Norwegian *Klebsiella pneumoniae* study group for collection of Norwegian isolates.

## Transparency declarations

The authors have no conflicts of interest to disclose.

MMCL, KEH and IHL conceived the study, performed data analyses and wrote the manuscript. KLW, RRW, AF, KEH and IHL contributed additional data analysis and interpretation. AF and IHL provided isolates. LMJ performed DNA extractions and Nanopore sequencing. All authors edited and approved the manuscript.

## Supplementary data

**Supplementary data 1:** strain table (csv)

**Supplementary data 2:** genome summary of Norwegian ST15 strains completed in this study (.docx)

**Supplementary figure. Coverage of mosaic plasmid pKp104014 amongst Norwegian and global ST15 genomes.**

Recombination-free, maximum likelihood phylogenetic tree (left) is as shown in **Fig. 1**, with tips coloured by the country of isolation (for n≥4 genomes) or geographical region as indicated. Heatmap indicates the presence (coloured; see legend in top right corner) or absence (white) of AMR genes and the KpVP-1 backbone annotated in plasmid pKp104014 in each genome, with key virulence genes labelled.

